# The association of MEG3 lncRNA with nuclear speckles in living cells

**DOI:** 10.1101/2022.05.11.491451

**Authors:** Sarah E. Hasenson, Ella Alkalay, Mohammad K. Atrash, Alon Boocholez, Julianna Gershbaum, Hodaya Hochberg-Laufer, Yaron Shav-Tal

## Abstract

Nuclear speckles are nuclear bodies containing RNA-binding proteins as well as RNAs including long non-coding RNAs (lncRNAs). MEG3 is a nuclear retained lncRNA that was identified to be associated with nuclear speckles. To understand the association dynamics of MEG3 lncRNA with nuclear speckles in living cells we generated a fluorescently-tagged MEG3 transcript that could be detected in real-time. Under regular conditions, transient association of MEG3 with nuclear speckles was observed, including a nucleoplasmic fraction. Conditions under which transcription or splicing were inactive, which are known to affect nuclear speckle structure, showed prominent and increased association of MEG3 lncRNA with the nuclear speckles, specifically forming a ring-like structure around the nuclear speckles. This contrasted with MALAT1 lncRNA that is normally highly associated with nuclear speckles, which was released and dispersed in the nucleoplasm. Under normal conditions MEG3 dynamically associated with the periphery of the nuclear speckles, but under transcription or splicing inhibition, MEG3 could also enter the center of the nuclear speckle. Altogether, using live-cell imaging approaches we find that MEG3 lncRNA is a transient resident of nuclear speckles and that its association with this nuclear body is modulated by the levels of transcription and splicing activities in the cell.

## 1. Introduction

The nucleus of higher eukaryotes contains various membraneless sub-nuclear compartments termed nuclear bodies [1,2]. One such body is the nuclear speckle which is host to many different types of RNA processing factors as well as long non-coding RNAs (lncRNAs) [3–5]. While most nuclear speckle components are splicing factors, many factors involved in the gene expression pathway can be found within. These include factors involved in transcription, RNA modifications and mRNA export regulation. Nuclear speckles were first identified by electron microcopy as 25-50 non-randomly distributed irregular shaped nuclear structures of varying sizes and were named inter-chromatin granule clusters (IGCs). The components of IGCs were later discovered to be factors involved mostly in pre-mRNA splicing [4,6]. Different sub-regions are identified within the nuclear speckles, as determined by super-resolution microscopy, and different components are detected in them [7]. For instance, proteins such as SRRM2 and SON that are known markers for nuclear speckles, can be found in the core region of the structure, while the splicing factor U2B and ncRNAs such as MALAT1 and snRNAs localize in the periphery of the nuclear speckles.

Nuclear speckles can take part in the regulation of gene expression but their exact roles are unclear [4]. It is suggested that nuclear speckles can function as storage and recycling sites for splicing factors [8,9]. Live-cell imaging studies have demonstrated splicing factors leaving the nuclear speckles supposedly to be recruited to transcription sites while FRAP analysis has shown that factors are constantly moving in and out these structures [10,11]. Moreover, both transcription inhibition and splicing inhibition lead to the accumulation of splicing factors in the nuclear speckles, which then grow in size and take on a rounder shape instead of the typical irregular structure, suggesting that under conditions with no splicing, the splicing factors only enter the nuclear speckles but do not leave [12,13]. We recently proposed that nuclear speckles can take part in buffering the levels of splicing factors in the nucleoplasm and as such regulate the amount splicing factors that are available for active splicing [14].

Nuclear speckles are also suggested to function as ‘gene expression hubs’ since they have been shown to have an enhancing effect on transcription and RNA processing activities for certain genes [15]. This may be influenced by the degree of proximity between the gene and the nuclear speckle, and by the recycling of the necessary factors between the gene and the nuclear speckles [11,16–20]. Interestingly, even though nuclear speckles may augment transcription of certain genes, they typically do not contain active genes nor DNA within, and the connection is probably determined by physical proximity.

Staining for poly(A) tails of transcripts has shown that nuclear speckles contain a substantial population of poly(A)+ RNAs [21–23]. No major change to this poly(A)+ RNA population was observed when transcription was blocked using α-amanitin, indicating that the poly(A)+ RNAs were not nascent mRNAs [22,24–26]. The lncRNA MALAT1 was the first poly(A)+ RNA identified as a key resident of nuclear speckles whereas the related lncRNA NEAT1 was found to localize in associated structures termed paraspeckles [27–30]. Most lncRNAs are nuclear retained and can also reside in sub-nuclear compartments [31]. MALAT1 is nuclear retained and its association with the nuclear speckles is transcription dependent and its primary function is alternative splicing regulation [32–34]. Many lncRNAs are transcribed by RNA polymerase ll and contain a 5’-cap and 3’ poly(A) tail. Generally, lncRNAs are expressed at lower levels than protein-coding mRNAs, and they are more tissue- or cell-type specific [31,35,36].

Extensive studies have revealed some lncRNAs functions, but the vast majority remains uncharacterized even at the most basic levels, such as subcellular localization and absolute abundance. In many cases, lncRNAs localize at their site of action, which therefore can provide insights into their biological function. Single molecule fluorescence *in situ* hybridization (smFISH) was used to identify the expression patterns of lncRNAs. One of these lncRNAs is Maternally expressed gene 3 (MEG3) [36,37]. This gene is located on chromosome 14 in human cells and its expression is regulated by differentially methylated promoters from the Dlk-MEG3 imprinting locus on the same chromosome [38–40]. MEG3 is expressed at high levels in the brain, placenta and endocrine glands [41–43]. It is involved in different pathways and is expressed in many diseases as well as having a tumor suppressor activity by selective activation of p53 response [44]. The underlying causes for loss of MEG3 expression in tumors are many, such as gene deletion, promoter hypermethylation and hypermethylation of the intergenic regions [45].

In this study we examined the dynamics of the MEG lncRNA in the nucleus of living cells. MEG3 is expressed in primary cells, while cancer cell lines usually express it at low levels. Therefore, we generated an inducible MEG3 gene that contains elements for tagging RNA in living cells, in a cancer cell line. We find that MEG3 is a nuclear retained lncRNA that can transiently associate with nuclear speckles under regular conditions, but when transcription or splicing inhibition are induced, the association with the periphery of nuclear speckles becomes predominant forming a distinct ring-like region surrounding the nuclear speckle.

## 2. Materials and Methods

### 2.1. Cell culture

U2OS Tet-On human osteosarcoma cells were maintained in low glucose Dulbecco’s modified Eagle’s medium (DMEM) (Biological Industries, Israel) containing 10% FBS (HyClone Laboratories). Stable expression of MEG3-MS2 + YFP-MS2 was obtained using PolyJet™ (SignaGen Laboratories) transfection, and selection with puromycin (1 μg/ml, Invivogen). Cells were grown at 37°C and 5% CO2. hFF (human foreskin) cells were maintained in high glucose DMEM (Biological Industries, Israel) containing 10% FBS. HepG2 (human liver cancer) cells were maintained in MEM-EAGLE Earle’s Salt Base medium (Biological Industries, Israel) containing 10% FBS. MEG3-MS2 transcription was induced by the addition of doxycycline (15 μg/ml) to the medium for 48 hrs.

For transcription inhibition, cells were grown on coverslips and incubated at 37°C for 2 hrs with either actinomycin D (5 μg/ml, Sigma) or DRB (50 μg/ml, Sigma) or 6 hrs with α-amanitin (30 μg/ml, Sigma) before fixation for 20 min in 4% PFA. To release from the DRB treatment, cells were washed with fresh medium for 30 min at 37°C before fixation. For splicing inhibition, cells were grown on coverslips and incubated at 37°C for 6 hrs with Pladienolide B (0.5 μM, Santa Cruz) before fixation for 20 min in 4% PFA. To release from the PLB treatment, cells were washed with fresh medium for 30 min at 37°C before fixation.

### 2.2. Plasmids and transfections

Cloning of MEG3 (NR_033358) under Tet-On control: A plasmid containing MEG3 was synthesized (Rhenium) [46] and cloned into the pTRE plasmid containing 24x MS2 repeats, by PCR amplifying the MEG3 sequence with primers matching the beginning and the end sequence with the addition of a BsrGI restriction site. The pTRE-24x-MS2 plasmid and the MEG3 PCR product were ligated following BSRGI restriction. The pTRE MEG3-MS2 plasmid was stably co-transfected with a YFP-MS2-A1 coat protein plasmid [47] into U2OS-Tet-On cells together with a plasmid for puromycin antibiotic selection. Different MEG3-MS2 cell clones were generated and used in the study. The truncated MEG3 Ex1-3-MS2 clone was created with: forward primer 5-AAGCTTTGTACAAGCCCCTAGCGC’-3’ matching the beginning of the sequence and primer 5’-TAAGGTGTACAGCTGATGCAAGGA-3’ matching the end of exon 3. Both primers contain the BsrGI restriction site for insertion into the pTRE-24x-MS2 plasmid. pTRE-MEG3 Ex1-3-MS2 and CFP-Clk1 were transfected using Polyjet (SignaGen Laboratories) according to the manufacturer’s instructions.

### 2.3. Immunofluorescence

Cells were grown on coverslips, washed with PBS and fixed for 20 min in 4% PFA. U2OS Tet-On cells were then permeabilized in 0.5% Triton X-100 for 2 min after which they were blocked in BSA. HFF and HepG2 were permeabilized twice in 0.1% Triton X-100 for 5 min. Cells were immunostained for 1 hr with a primary antibody, and after subsequent washes the cells were incubated for 45 min with secondary fluorescent antibodies. Primary antibodies: Mouse anti-SC35 (which marks SRRM2 [48]), rabbit anti-SON (Sigma), rabbit anti-hnRNPK (Abcam) and rabbit anti-SRSF7 (Santa Cruz). Secondary antibodies: Alexa647-labeled goat anti-mouse IgG, Alexa594-labeled goat anti-mouse IgG, Alexa647-labeled donkey anti-rabbit IgG (Life Technologies), dyLight488-labeled goat anti-rabbit IgG and Alexa488-labeled goat anti-mouse IgG (Abcam). The cells were mounted in mounting medium.

### 2.4. Fluorescence microscopy

Wide-field fluorescence images were obtained using the CellSens system based on an Olympus IX81 fully motorized inverted microscope (60x Planpon O objective 1.42 [NA]) fitted with an Orca-Flash4.0 #2 camera (Hamamatsu, Bridgewater, NJ), rapid wavelength switching, and driven by the CellSens software; or the CellR system based on an Olympus IX81 fully motorized inverted microscope (60x PlanApo objective 1.42 [NA]) fitted with an Orca-AG CCD camera (Hamamatsu, Bridgewater, NJ), rapid wavelength switching, and driven by the CellR software. For time-lapse imaging, cells were plated on glass-bottomed tissue culture plates (MatTek, Ashland, MA) in DMEM medium containing 10% fetal calf serum at 37°C. The microscope is equipped with an on-scope incubator which includes temperature and CO2 control (Life Imaging Services, Reinach, Switzerland). For long-term imaging, several cell positions were chosen and recorded by a motorized stage (Scan IM; Märzhäuser, Wetzlar-Steindorf, Germany). Cells were typically imaged in three dimensions (3D) (11 Z planes per time point) every 5 or 10 minutes. For quantifications of MEG3 intensity in association with the nuclear speckle, the cells were imaged in 31 Z planes per time point. Rapid live-cell imaging was performed by imaging every 300 msec for 60 sec in one Z plane. Live-cell imaging and 3D stacks were deconvolved using Huygens (Scientific Volume Imaging) and analyzed by Imaris (Oxford Instruments). The perimeter of the of the nuclear speckle area analyzed with the Imaris extended outward with a threshold value of 0.25 or 0.25 μm from the surface of the nuclear speckles. Colocalization analysis of two channels was performed using an ImageJ Macro.

Super resolution fluorescence and confocal images were obtained using the Leica SP8 STED confocal microscope (63x HC PL APO objective 1.40 [NA]). For STED imaging, two continuous wave lasers at 592 nm and 660 nm were used, which allow the use of fluorophores up to 595 nm in super-resolution mode. The white light laser (WLL) function was used for the confocal imaging.

FRAP experiments were performed using a Leica TIRF microscope (63X HC PL APO objective 1.40 [NA]), as previously described [14]. Before and after the bleaching, cells were imaged in the YFP channel for the detection of YFP-MS2 (MEG3) and in the CFP channel for the detection of Cer-SRSF3 (nuclear speckle factor). MEG3 signal at a specific nuclear speckle was photobleached using the 553 nm laser. Six pre-bleach images were acquired. Post-bleach images were acquired in a sequence of 3 time frequencies: 15 images every 3 sec, 15 images every 6 sec, and 26 images every 30 sec. For each time-point, the background taken from a region of interest (ROI) outside of the cell was subtracted from all other measurements. T(t) and I(t) were measured for each time-point as the average intensity of the nucleus and the average intensity in the bleached ROI, respectively. The average of the pre-bleach images used as the initial conditions are referred to as Ti = nuclear intensity and Ii = intensity in ROI before bleaching. Ic(t) is the corrected intensity of the bleached ROI at time t:

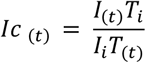

The diffusion of the YFP-MS2 protein during the FRAP analysis was disregarded since the diffusion rate of free nucleoplasmic YFP-MS2 is very rapid, while the bound YFP-MS2 is associated with high affinity to the RNA, and does not detach or diffuse [49,50].

### 2.5. RNA FISH

Cells were seeded on 18 mm coverslips and fixed for 20 min in 4% paraformaldehyde (PFA), and then permeabilized twice in 0.1% Triton X-100 for 5 min. Coverslips were then washed twice with 10% formamide for Stellaris probes or 15% formamide for FLAP [51] probes diluted in 4xSSC. For smFISH on endogenous transcripts, fluorescent-labeled DNA probes that target the MEG3 sequence (570 nm, ~10 ng probe, Stellaris; or 670 nm, ~5 ng probe, FLAP), MALAT1 sequence (570 nm, ~10 ng probe, Stellaris) or NEAT1 sequence (570 nm, ~10 ng probe, Stellaris) were hybridized overnight at 37°C in a dark chamber in 10% formamide or 15% formamide, respectively. The next day cells were washed twice with 10% formamide for Stellaris probes or 15% formamide for FLAP probes diluted in x4 SSC for 30 min at 37°C and then washed with 1x PBS. The coverslips were mounted with mounting medium.

For MS2 RNA FISH, cells were left overnight in 70% ethanol at 4°C and then fluorescently labeled probes that target the MS2 repeats (670 or 570, ~10 ng probe per coverslip) were hybridized overnight at 37°C in a dark chamber in 40% formamide. The next day, cells were washed twice with 40% formamide diluted in SSC x4 for 15 min followed by two additional washes with 1X PBS for one hr each. Coverslips were mounted with mounting medium.

### 2.6. siRNA knockdowns

Cells were transfected with siRNA Son (Thermo Scientific), siRNA hnRNPK (IDT), siRNA SRSF7 (Invitrogen) or a negative scrambled control (IDT), using Lipofectamine 2000.

### 2.7. FACS cell cycle analysis

10^6^ wildtype or MEG3 U2OS Tet-On cells were seeded in a 10 cm plate. Doxycycline (15 μg/ml) was added to the medium for 48 hrs. Trypsin was used to detach cells from plates. Cells were washed with PBS and then fixed in PFA 4% for 15 min. Cells were washed and resuspended in DAPI 1 mg/ml in PBS and 10% Triton x-100 at RT for 8 min. Cells were washed the pellet was resuspended with 1 ml PBS and analyzed in a Flow Cytometer LSRFortessa (BD Biosciences, NJ, USA). Analysis of the data output was preformed via FlowJo v10.8 software using Watson model to calculate the percent of cells in each stage of the cell cycle.

### 2.8. Statistical analysis

Three-parameter asymptotic regression was used to analyze FRAP experiments. Each replicate was fit to a curve defined by the equation Y=a-(a-b) e˄(-cX) where X is time and Y is the relative intensity, a (plateau) is the maximum attainable intensity, b (init) is the initial Y value (at time=0) and c (m) is proportional to the relative rate of increase for intensity when time increases. The regression fitting was performed using the drm function from the drc R package and DRC.asymReg self-starting function from the aomisc R package. Each parameter (init, m and plateau) was then compared between all treatments with a one-way nested ANOVA, followed by pairwise comparisons of mean values between treatments. Finally, p-values were adjusted for multiple comparisons with the Benjamini-Hochberg (FDR) procedure. The function lmer from lmerTest package, and emmeans from emmeans package were used.

## 3. Results

### 3.1. Detection of MEG3 lncRNA in living cells

To facilitate the detection of MEG3 lncRNA in living cells, we generated an osteosarcoma U2OS cell line in which an inducible MEG3 gene was stably expressed. Most cancerous cell lines express MEG3 lncRNA at very low levels and so osteosarcoma cells do not have background levels of MEG3 [52–54]. The gene sequence contains a series of 24x MS2 sequence repeats at the 3’-end, used to facilitate tagging of RNA in living cells [55,56]. The expression of MEG3-MS2 was induced under the Tet-On promotor system following the addition of doxycycline (Dox) to the cell medium (Figure 1A). The MEG3-MS2 gene was co-transfected with the YFP-MS2 coat protein (YFP-MS2-CP) that specifically binds to the repeated stem-loop structures in the 3’-end of the transcript formed by the 24x MS2 repeats, thus allowing for detection of the RNA in living cells (Figure 1B). Single transcripts (lncRNPs) could be detected in the nucleoplasm of the dox-induced U2OS cells, and were excluded from nucleloli. The active site of transcription of the MEG3-MS2 gene was also observed. Expression of MEG3-MS2 did not affect the cell cycle of U2OS cells (Figure S1A). The distribution patterns of MEG3-MS2 lncRNA in U2OS cells were predominantly nuclear and similar to those of the endogenous MEG3 transcript as observed in Hepatocellular carcinoma (HepG2) cells, a cancerous cell line which does express MEG3 [57]. The endogenous MEG3 lncRNA was detected using RNA FISH probes to MEG3 (Figure S1B).

**Figure 1.**
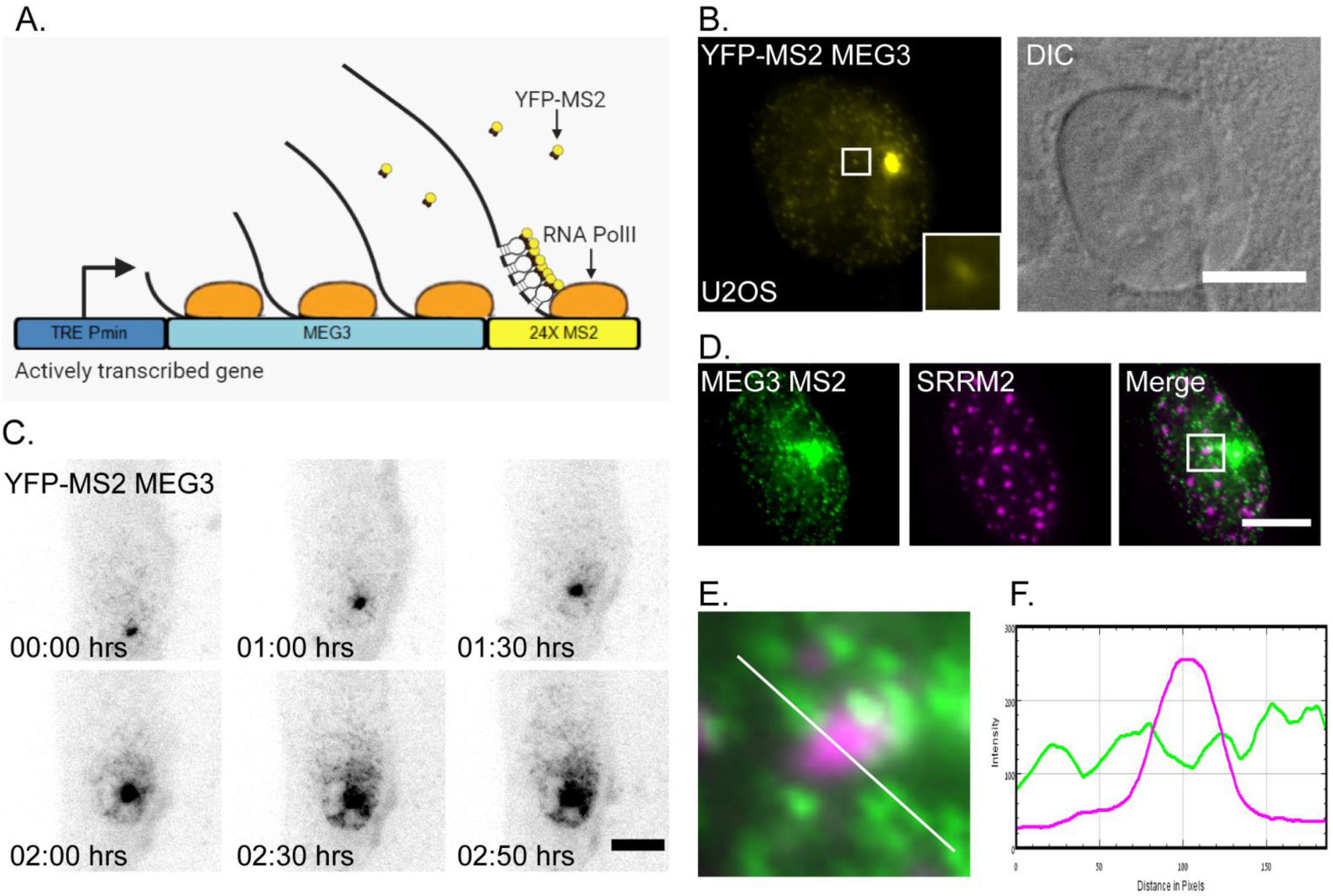
MEG3-MS2 expressed in U2OS cells partially associates with the nuclear speckles. (**A**) Schematic depiction demonstrating the Tet-On MEG3-MS2 expression system. (**B**) U2OS cells stably expressing MEG3-MS2 tagged with YFP-MS2-CP after Dox activation (yellow) and DIC (grey). Small dots are MEG lncRNPs and the large dot is the active gene. Enlargement of an area with transcripts in the boxed region. Bar= 10 μm. (**C**) Frames from a live-cell movie (**Movie S1**) of the transcriptional induction (dox) of MEG3-MS2 expression. The grey image is an inverted version of the transcript tagged with YFP-MS2-CP (black). Images were captured every 10 min for the duration of 3 hrs. Bar= 5 μm. (**D**) MEG3-MS2 tagged with YFP-MS2-CP (green) is partially associated with nuclear speckles stained with an antibody to the SRRM2 nuclear speckle marker (magenta). (**E**) Enlargement of MEG3 and SRRM2 and (**F**) intensity line analysis from the boxed region in D along the white line. Bar= 10 μm.

There was no cytoplasmic MEG3 signal detected in the U2OS cells expressing MEG3-MS2, demonstrating that MEG3 is a nuclear retained lncRNA, as previously documented in normal fibroblasts such as hFF (human foreskin fibroblasts), S27 (Human Foreskin Fibroblast), W138 (Human Lung Fibroblast) cells as well as a cancerous cell line such as hLF (Human Lung Fibroblast Epidermoid Carcinoma) [36,41]. A live-cell movie of MEG3-MS2 tagged with YFP-MS2-CP showed the gradual increase in MEG3 transcription and the distribution of the lncRNPs in the nucleoplasm over a 3-hour period (Figure 1B and C, Movie S1). Active transcription was observed about 15 minutes after Dox induction and the lncRNPs began to spread in the nucleoplasm at around 1.5 hours and onwards. The transcripts were not evenly distributed throughout the nucleoplasm. We therefore examined whether MEG3-MS2 might be associated with nuclear speckles as previously suggested [36], like the two other nuclear retained lncRNAs, MALAT1 and NEAT1, which are known to be in association with these nuclear bodies [27]. It was previously shown that lncRNAs and proteins localize within different regions of the nuclear speckle structure. Some proteins are found at the center of the nuclear speckle while the MALAT1 lncRNA is found at the periphery [7]. Some of the MEG3 lncRNA signal was in close proximity with the SRRM2 protein, a core nuclear speckle protein (Figure 1D-F). Altogether, there appears to be a partial level of association between the nuclear speckles and the MEG3 signal.

### 3.2. MEG3 localizes to nuclear speckles upon transcription inhibition

RNA FISH to the MALAT1 lncRNA shows a clear overlap in the position of the lncRNAs and nuclear speckles (Figure 2A), as expected [27]. It was previously shown that MALAT1 and MEG3 colocalize to a high extent in hFF cells [36], and therefore the association of MEG3 with nuclear speckles is expected, although the MEG3 signal appears to be more dispersed compared to the MALAT1 signal which is more concentrated within the nuclear speckles. To further examine the association of MEG3 with nuclear speckles, we used transcription inhibition conditions. The latter treatment causes nuclear speckles to retain an excess of splicing factors that are no longer needed in the nucleoplasm, leading to the rounding up and enlargement of the nuclear speckles, compared to regular conditions [11,58]. Transcription inhibition was induced by treating cells with 5,6-dichloro-1-β-D-ribofuranosylbenzimidazole (DRB) that inhibits transcription elongation [59]. While MALAT1 normally resides in the nuclear speckles, transcription inhibition by DRB leads to its dispersal in the nucleoplasm, whereas nuclear speckles remain intact [32] (Figure 2A). Similarly, the NEAT1 lncRNA known to localize to paraspeckles under regular conditions, also disperses after treatment with DRB [60] (Figure 2B). Interestingly, for MEG3 lncRNA, the opposite was observed. Transcription inhibition with DRB led to close localization of the MEG3 lncRNAs to the nuclear speckles and a ring-like pattern appeared at the nuclear speckle periphery (Figure 2C). The rounding up of the nuclear speckles could be observed in the DRB-treated cells, as expected. The redistribution of MEG3 in DRB-treated cells was then followed using live-cell imaging after 15 minutes of treatment. The recruitment of MEG3 to the nuclear speckle began after 35 minutes of DRB treatment (20 min into the movie) and reached full development by 1 hour (Figure 2D and Movie S2).

**Figure 2.**
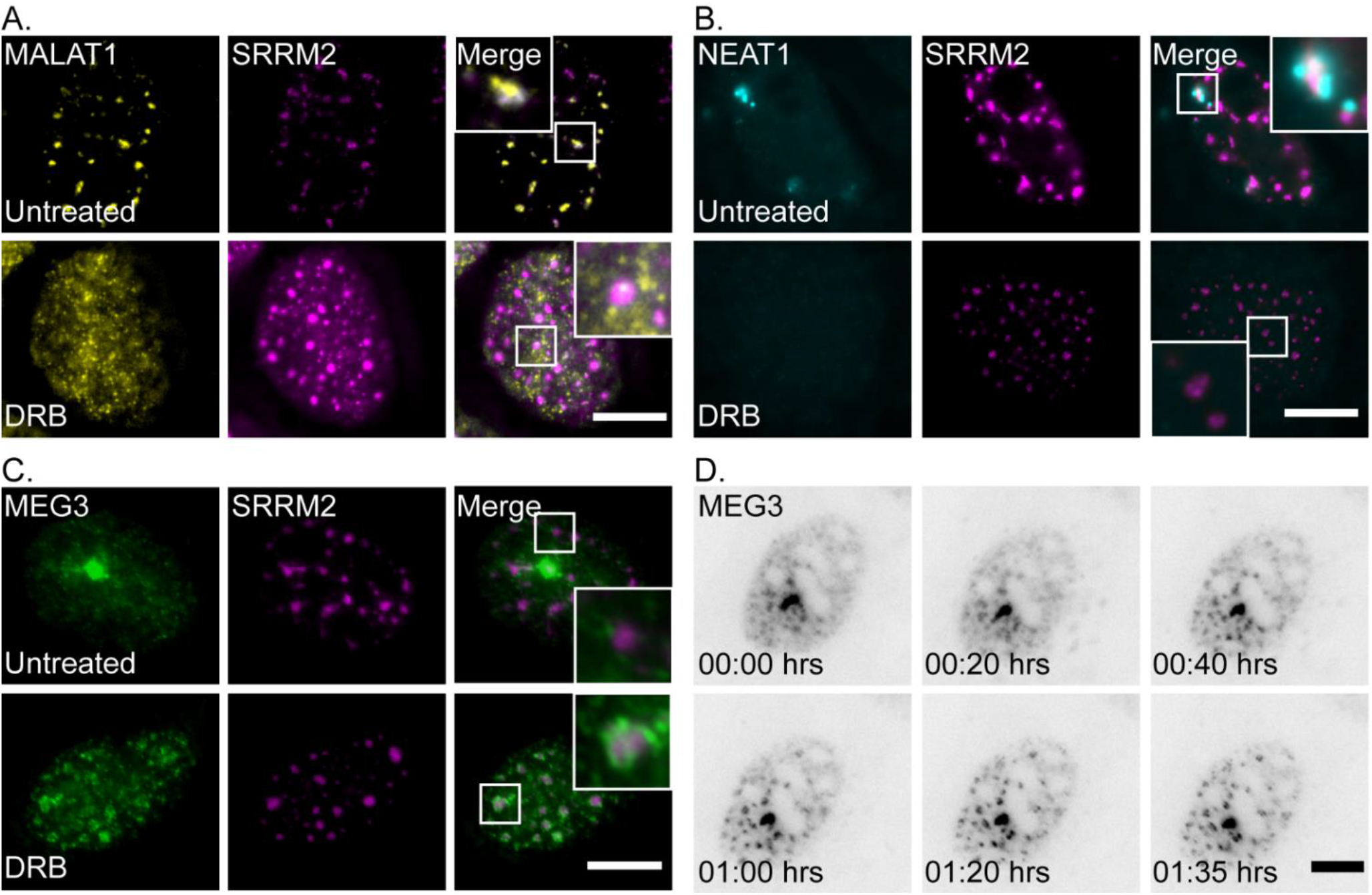
MALAT1 and NEAT1 lose nuclear speckle association in response to transcription inhibition while MEG3 is recruited to the nuclear speckle periphery. (**A**) (Top) RNA FISH to endogenous MALAT1 lncRNA (yellow) overlapped with antibody staining for nuclear speckles marked with SRRM2 (magenta) in untreated cells. (Bottom) When treated with the transcription inhibitor DRB (2 hrs), MALAT1 lncRNA dispersed in the nucleoplasm and the overlap with SRRM2 was lost. Enlargements of designated areas are in the boxed regions. (**B**) (Top) RNA FISH to endogenous NEAT1 (cyan) overlapped with the periphery of nuclear speckles marked with an SRRM2 (magenta) in untreated cells. (Bottom) In DRB-treated cells, NEAT1 dispersed in the nucleus. (**C**) (Top) MEG3-MS2 tagged with YFP-MS2-CP (green) partially associated with nuclear speckles marked with an antibody to SRRM2 (magenta) in untreated cells. (Bottom) When treated with DRB, MEG3 localized to the nuclear speckles and there was increased association with the SRRM2 signal. Bar= 10 μm. (**D**) Frames from a live-cell movie (**Movie S2**) of a dox-induced MEG3-MS2 cell expressing the transcript coated with YFP-MS2-CP, starting 15 after DRB treatment. Images were captured every 5 min for the duration of 1.5 hrs. The grey image is an inverted version of the transcript tagged with YFP-MS2-CP (black). Bar= 5 μm.

Different types of transcription inhibitors affect distinct steps of the transcription process. We also tested actinomycin D (ActD) that inhibits the access to DNA for all mammalian RNA polymerases, and α-amanitin that specifically but slowly inhibits RNA polymerase II. Both ActD and α-amanitin had similar effects on MEG3 distribution, compared to DRB (Figure 3A and S2).

**Figure 3.**
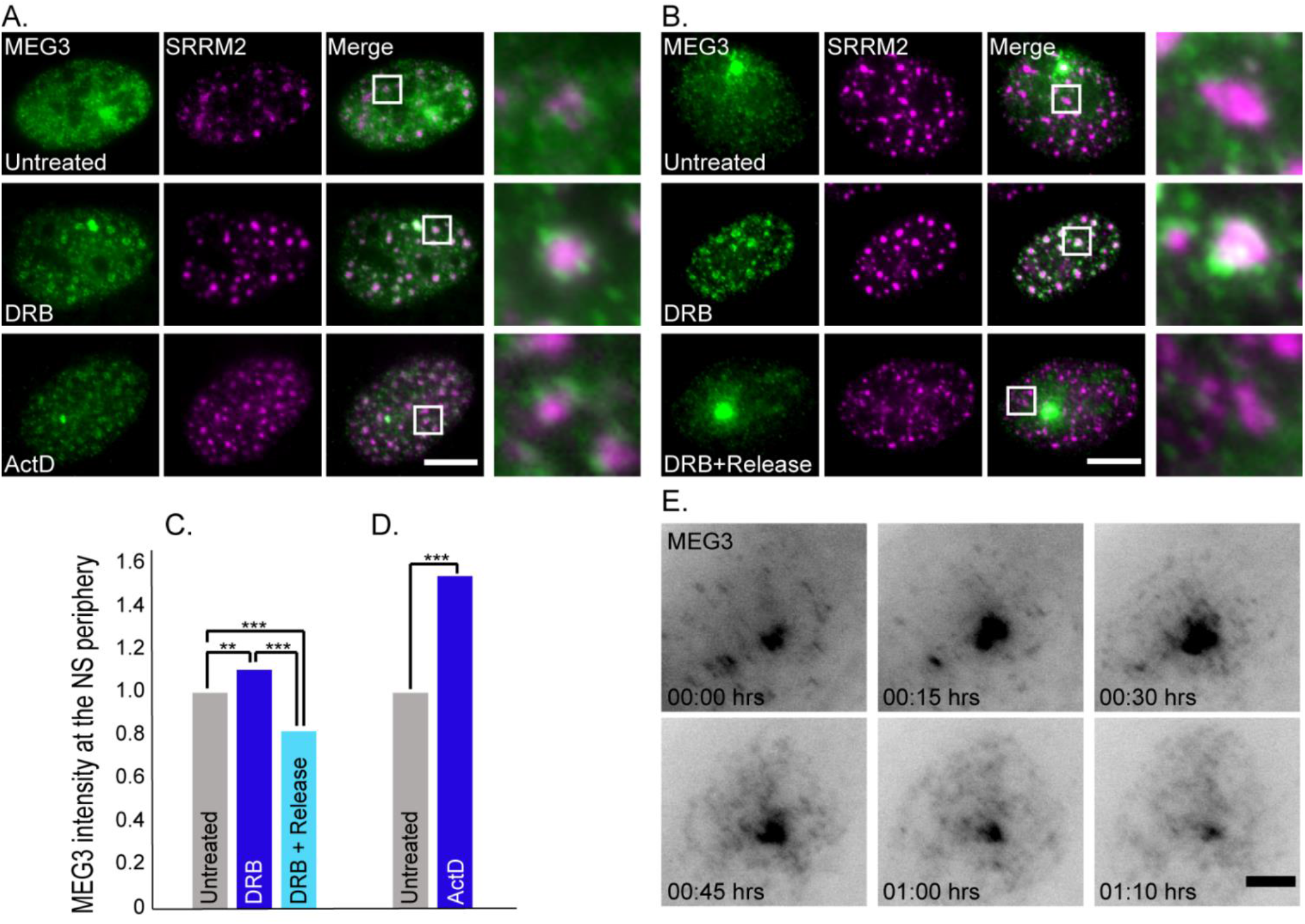
The increase in MEG3-nuclear speckle association in response to transcription inhibition is reversible. (**A**) MEG3 YFP-MS2-CP tagged transcripts (green) and antibody staining of SRRM2 (magenta) under untreated or different transcription inhibition treatments (2 hrs of DRB or ActD before fixation). Enlargements of the boxed regions appear on the right. (**B**) RNA FISH of MEG3-MS2 transcripts with fluorescent probes to the MS2 region of the lncRNA (green) and antibody staining of SRRM2 (purple). (Top) Untreated cells, (middle) DRB-treated cells (2 hrs) and (bottom) cells treated with DRB for 2 hrs and then released in medium for 30 min before fixation. Bar= 10 μm. (**C**) Measurements of the changes in MEG3 intensity at the nuclear speckle (NS) periphery under untreated conditions (#cells=94, #NS=1165), DRB treatment (#cells=61, #NS=1031), and after 30 minutes release from DRB (#cells=108, #NS=1331) (**p<0.001, ***p<0.0001). (**D**) Measurements of the changes in MEG3 association with nuclear speckles under untreated conditions (#cells=124, #NS=1300) and ActD treatment (#cells=99, #NS=1024) (***p<0.0001). (**E**) Frames from a live-cell movie (**Movie S3**) of stably expressing MEG3-MS2 YFP-MS2-CP after release from DRB. Images were captured every 5 min for the duration of 1.25 hrs. The grey image is an inverted version of the transcript tagged with YFP-MS2-CP (black). Bar= 5 μm.

The association of protein factors with nuclear speckles is highly dynamic, whereby they are continuously moving in and out of these structures [11,61]. To test whether the MEG3 nuclear speckle localization was reversible, cells were treated with DRB (2 hrs), and then returned to regular medium for 30 minutes before fixation (Figure 3B). MEG3 localization to the nuclear speckles caused by transcription inhibition, dispersed after DRB removal. Still, the regular fraction of MEG3 associated with nuclear speckles was observed after release. Quantifications of the MEG3 intensity at the nuclear speckle periphery showed a significant increase both in DRB and in ActD-treated cells compared to untreated or DRB-released cells (Figure 3C and D). Live-cell imaging of cells treated with DRB for 2 hours and then released with medium, showed that the reversal of nuclear speckle localization was quite rapid and started 15 minutes after DRB removal (Figure 3E, S3 and Movie S3). The release from DRB was apparent by the return to the typical irregular shape of nuclear speckles.

We verified that the relocalization of MEG3 lncRNA to nuclear speckles also occurs for the endogenous transcript in HepG2 and hFF cells (Figure 4A). These cells were treated with DRB, and endogenous MEG3 transcripts were detected by RNA FISH. In both cell lines there was some MEG3 association with nuclear speckles under regular growth conditions, as seen in U2OS cells. Increased localization to the nuclear speckle periphery was seen upon DRB treatment (Figure 4B).

**Figure 4.**
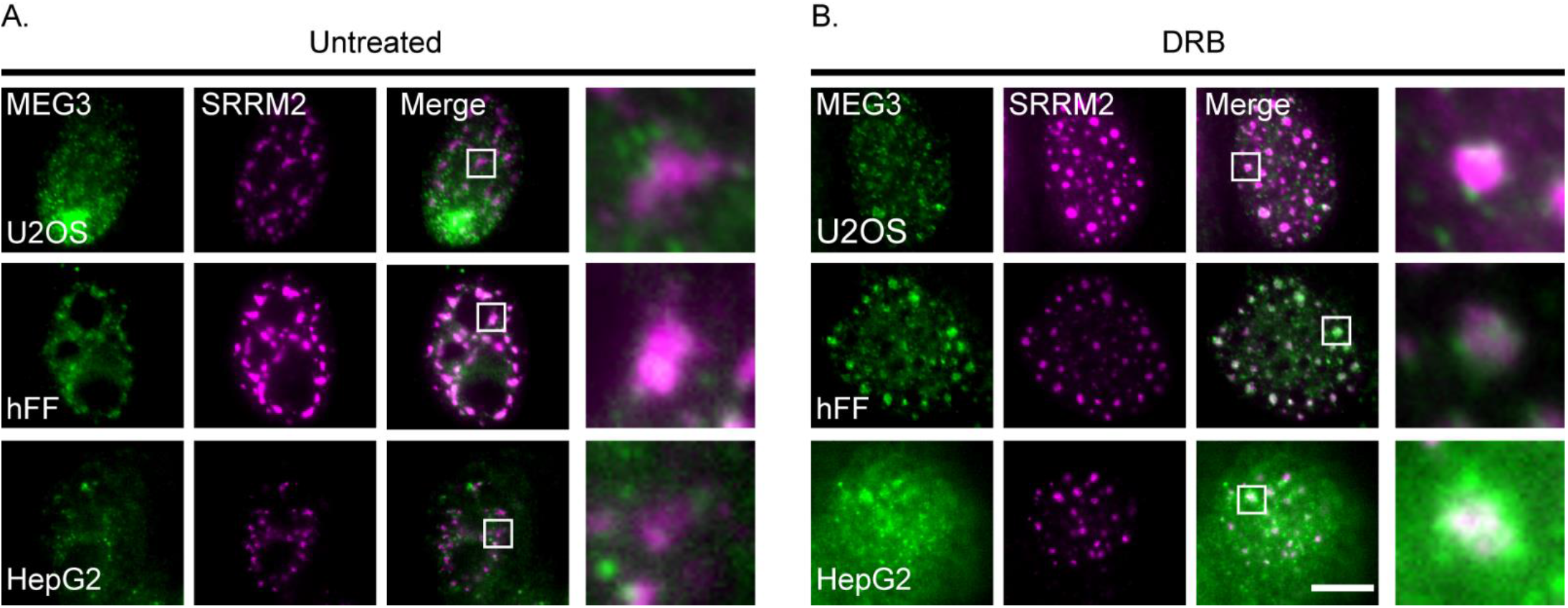
Endogenous MEG3 localizes to the nuclear speckle. (**A**) RNA FISH of MEG3 transcripts (green) and antibody staining to SRRM2 (magenta) in (Top) untreated U2OS MEG3-MS2 cells; (Middle) hFFs; and (Bottom) HepG2 cells. Enlargements of boxed areas are on the right. (**B**) Same as A with DRB treatment (2 hrs). Bar= 10 μm.

### 3.3. MEG3 localizes to nuclear speckles during splicing inhibition

Nuclear speckles contain a large variety of splicing factors. The phenomenon of rounding up of these structures during transcription inhibition is attributed to the return of splicing factors from the nucleoplasm to the nuclear speckles for recycling without the need to return to genes, since there are no active genes producing pre-mRNAs during transcription inhibition [11,58,62]. Splicing inhibition on its own has the same effect on nuclear speckle structure [63]. We therefore wished to examine whether splicing inhibition per se, without transcription inhibition, can affect the distribution of MEG3 lncRNA. Pladienolide B (PLB) is a compound that inhibits splicing [64] and indeed when the cells were treated with PLB, MEG3 transcripts formed the circular formations surrounding the nuclear speckles, resembling those observed during transcription inhibition. PLB treatment was also reversible but appeared to have somewhat slower dynamics than the release from DRB (Figure 5A, B).

**Figure 5.**
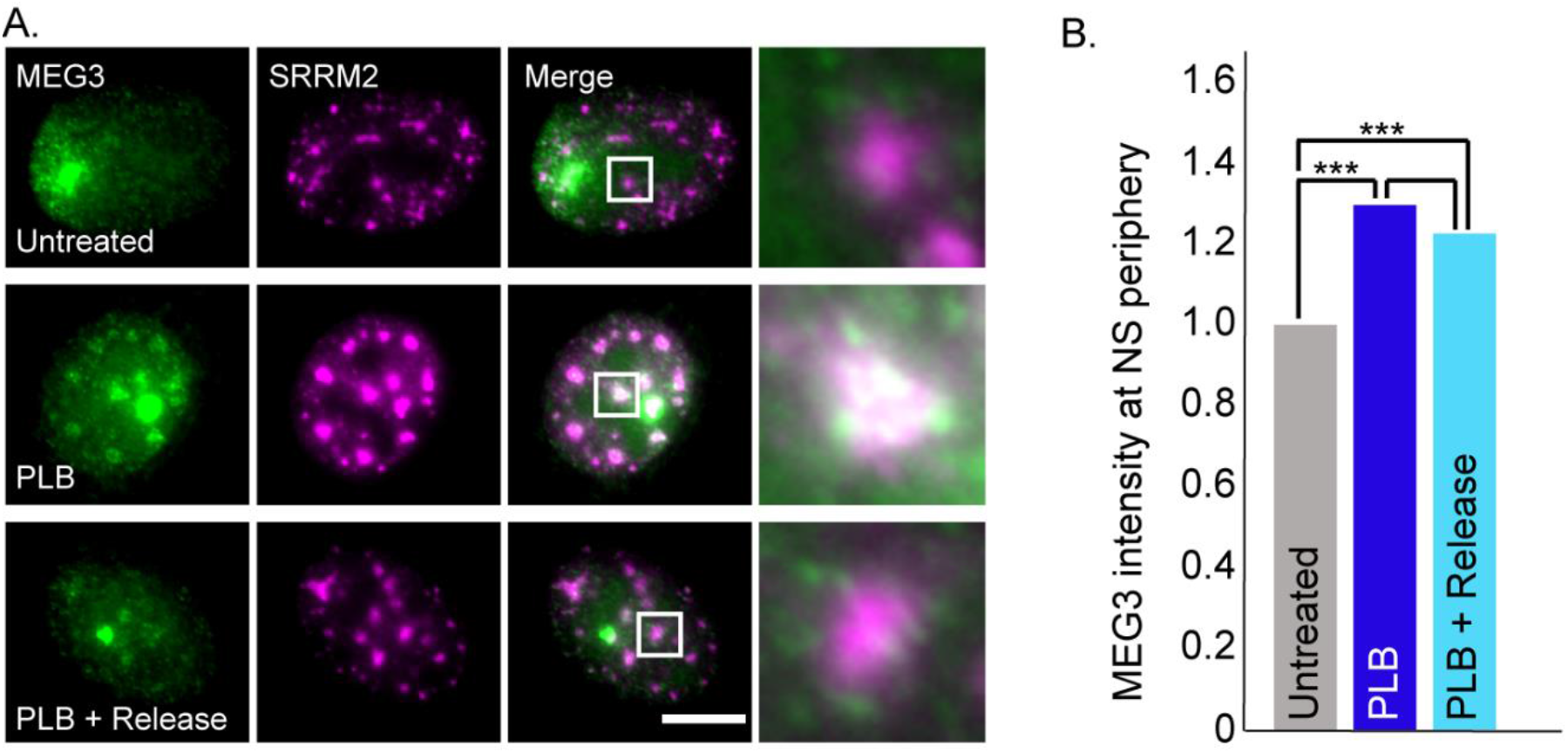
MEG3 surrounds nuclear speckles in response to splicing inhibition. (**A**) RNA FISH of MEG3-MS2 transcripts (green) and antibody staining of SRRM2 in nuclear speckles (purple). (Top) Untreated control cells, (middle) cells treated with PLB for 6 hrs and (bottom) cells treated with PLB for 6 hrs and then released 30 min before fixation. Bar= 10 μm. Enlargements of the boxed areas are on the right. (**B**) Plot demonstrating the changes in MEG3 association with nuclear speckles (NS) under untreated conditions (#cells=84, #NS =1057), PLB treatment (#cells=111, #NS=1212) and after a 30 min release from PLB (#cells=116, #NS=1403) (***p<0.0001).

### 3.4. The first 3 exons MEG3 sequences are responsible for the nuclear speckle-MEG3 interaction

The human *MEG3* gene has 8 exons. To examine which sequences in the lncRNA are connected to the localization to nuclear speckles a shorter clone of MEG3-MS2 was created. We decided to remove roughly half of the MEG3 transcript resulting in a sequence containing exons 1-3 (exon 3 is long compared to the other exons). The transcript was expressed in U2OS cells that were then treated with DRB. Transcripts containing exons 1-3 still localized to the nuclear speckles, indicating that the last five exons were not required for this localization (Figure 6A).

**Figure 6.**
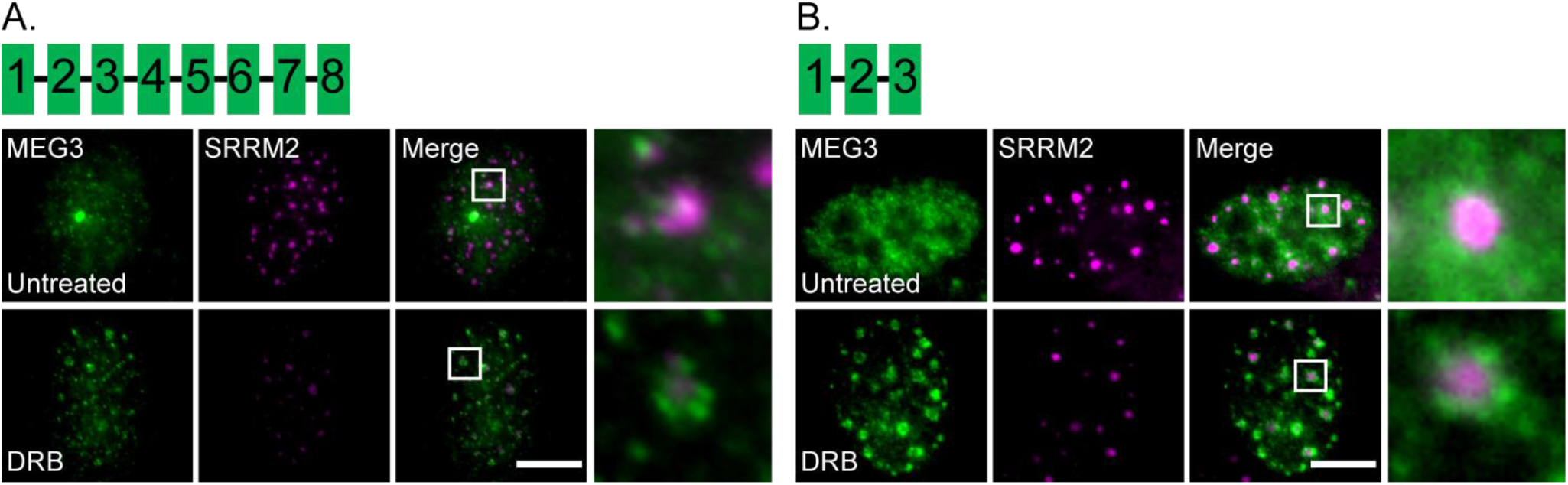
The last 5 exons of MEG3 are not needed for MEG3 localization around nuclear speckles during transcription inhibition. (**A**) RNA FISH of full length MEG3-MS2 and (**B**) truncated transcripts (exons 1-3; green) in U2OS cells with SRRM2 staining (magenta). The localization of the transcripts was examined in untreated control cells (top) and cells treated with DRB (2 hrs, bottom). Depictions of the MEG3 exons (green boxes) in each construct are shown above the images. Enlargements of the boxed areas are on the right. Bar =10 μm.

As demonstrated above, MEG3 repositioning in relation to the nuclear speckles can take place due to changes in the nuclear speckle structure occurring during transcription and splicing inhibition. Next, we knocked down different RNA-binding factors by use of siRNA, to see if we could affect MEG3 nuclear speckle localization. SRRM2 was used as a nuclear speckle marker to show that the nuclear bodies were still intact even though a specific protein was knocked down. The core proteins SON and SRRM2 appear to be essential for proper nuclear speckle formation, since their depletion can lead to different degrees of nuclear speckle disassembly or restructuring [48]. When SON was knocked-down and cells were transcriptionally inhibited, SRRM2 staining showed partial loss of nuclear speckle formation, however, MEG3 still appeared to localize to the nuclear speckle surrounding the remaining assemblies of SRRM2. This would suggest that SON is not responsible for the increased speckle localization during transcription/splicing inhibition (Figure 7). Knockdown of the SRSF7 splicing factor or of Heterogeneous nuclear ribonucleoprotein K (hnRNPK) that is known to bind lncRNAs [65] did not have any impact on MEG3 localization either (Figure S4A and B).

**Figure 7.**
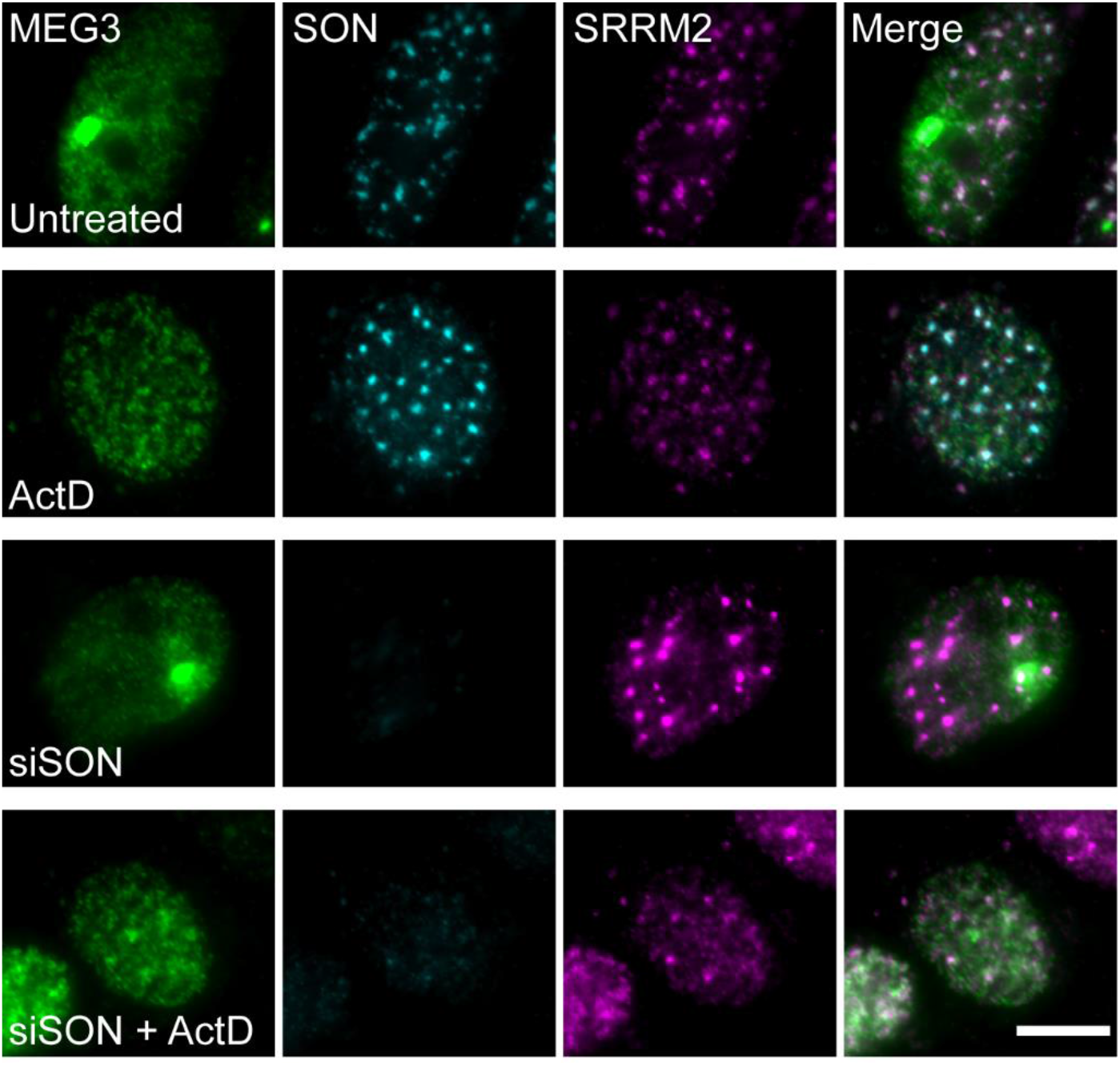
Depletion of SON protein does not change MEG3 association with nuclear speckles. Knockdown of SON (cyan) in cells labeled by RNA FISH for MEG3-MS2 transcripts (green) and staining SRRM2 (magenta). (Top) Untreated control cells, (top middle) cells treated with ActD for 2 hrs, (bottom middle) siRNA knockdown, (bottom) siRNA knockdown with ActD treatment for 2 hrs. Bar= 10 μm.

### 3.5. MEG3 dynamics at the nuclear speckle during transcription inhibition

We hypothesized that MEG3 once located at the nuclear speckles will remain there as long as the cell is under transcription inhibition conditions. This could be examined by fluorescence recovery after photobleaching (FRAP). Measuring the dynamics of a lncRNA at the nuclear speckle under transcription inhibition has not been examined so far since other lncRNAs such as MALAT1 and NEAT1 are released from the nuclear speckle under transcription inhibition conditions [29,32]. FRAP analysis was performed on cells where MEG3 was tagged with YFP-MS2-CP and co-expressed with Cerulean-SRSF3 as a nuclear speckle marker. MEG3 signal was photobleached and the return of the YFP signal was measured at the nuclear speckles in untreated and DRB-treated cells. MEG3 signal at the nuclear speckles in untreated cells showed ~90% recovery. In treated cells the recovery was slower and gradual, indicating increased association of MEG3 with the nuclear speckles (Figure 8A, B and Movie S4). Still, the signal did finally recover indicating that there is a certain degree of MEG3 exchange from the nuclear speckle over time. Next, we further examined this interchange of MEG3 transcripts between the nuclear speckle and the nucleus by tracking of the transcripts. MEG3 movement in relation to the nuclear speckle labeled with CFP-SRSF3 was followed and characterized using rapid live-cell imaging. In transcriptionally inhibited cells, MEG3 movement was confined to the nuclear speckles whereas in untreated cells the RNA had significantly larger displacement lengths. The limited movements after DRB treatment at the nuclear speckles showed that the transcripts moved back and forth at the nuclear speckle periphery, but did not tend to leave its vicinity (Figure 8C and D).

**Figure 8.**
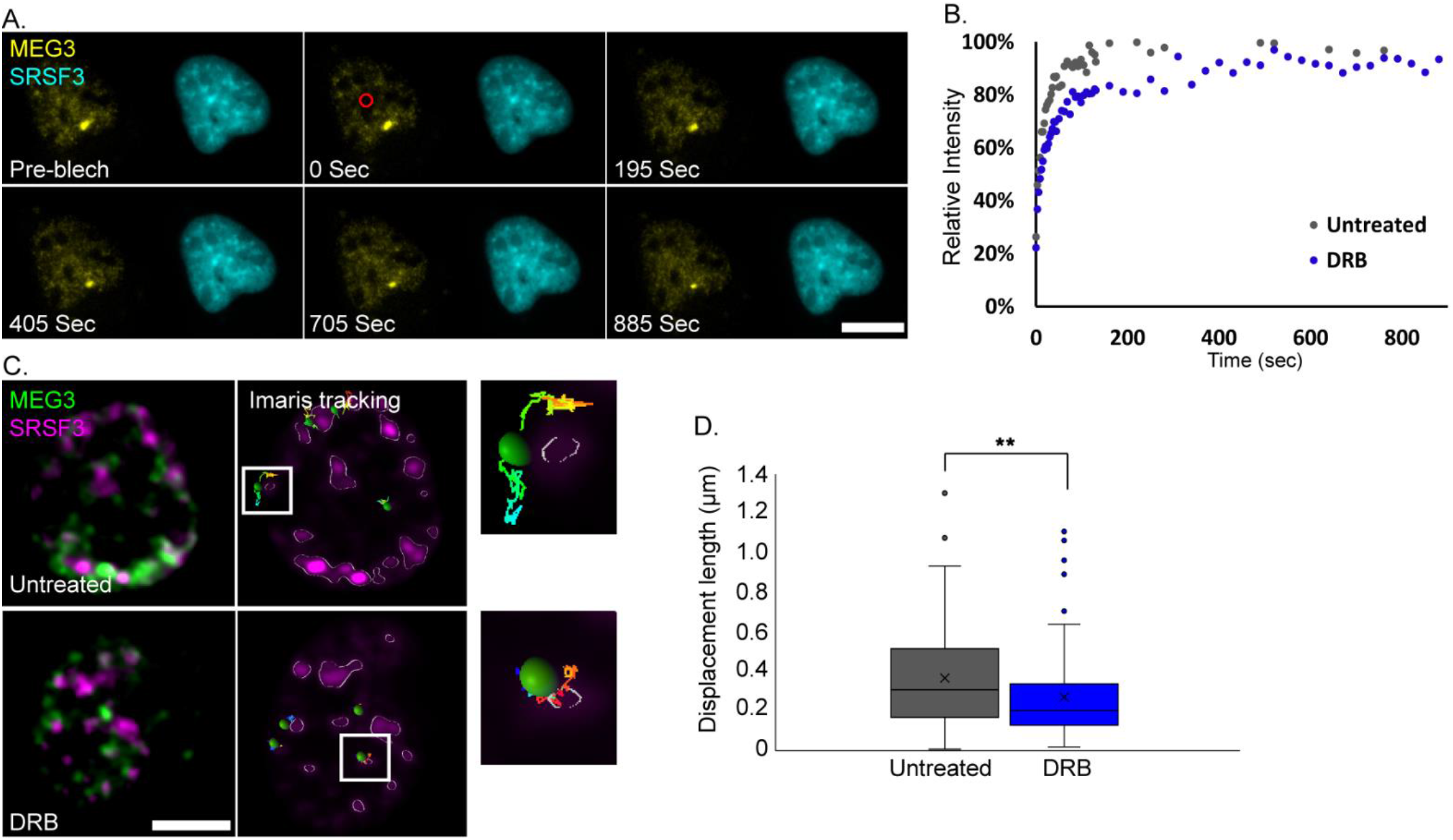
MEG3 is highly associated with nuclear speckles during transcription inhibition. (**A**) Frames from a FRAP experiment showing MEG3 YFP-MS2-CP signal (yellow), Cerulean-SRSF3 (cyan) in the same cell. Photobleached area - red circle. (**B**) FRAP recovery curves of MEG3 at nuclear speckles (#untreated cells cells=27, #DRB treated cells=21, *p<0.05 in the middle phase of the recovery curve) in untreated (gray) cells and DRB-treated (blue) cells. DRB treated cells were imaged during the second hour of the treatment. (**C**) Tracking MEG3 YFP-MS2-CP (green) in relation to the nuclear speckle marker Cer-SRSF3 (magenta). Unprocessed image (left) and the Imaris rendering (right) of untreated (top) and DRB treated (bottom) cells. Enlargements of frames from the boxed areas appear in the images. Bar= 10 μm. (**D**) Plot demonstrating the displacement length of MEG3 movements within a 0.25μm radius from nuclear speckles (NS) under untreated conditions (#cells=27, #RNA=108) and DRB treatment (#cells=25, #RNA=105) (**p<0.005).

Super-resolution microscopy has shown that nuclear speckles are layered structures and that the SRRM2 protein is most often found at the core of the nuclear body [7]. To examine the precise position of MEG3 lncRNA at the nuclear speckle and the changes occurring after transcription or splicing inhibition, we stained the core nuclear speckle factors SON and SRRM2, after which the cells were imaged using STED super-resolution microscopy. Before transcription inhibition, MEG3 lncRNA was loosely associated with the periphery of the nuclear speckle core, the latter harboring the SRRM2 and SON proteins. After treatment with DRB or ActD, prominent MEG3 signal was seen around the nuclear speckles. Interestingly, some MEG3 signal was also detected within the inner part of the nuclear speckle (Figure 9A and B). Still, the RNA and protein signals remained spatially separate even in the core of the nuclear speckle. Confocal imaging of HepG2 cells showed the endogenous MEG3 behaving in a similar manner. In untreated cells, MEG3 associated with the periphery of the nuclear speckles but after treatment with DRB, MEG3 localized to its core (Figure 9C and D).

**Figure 9.**
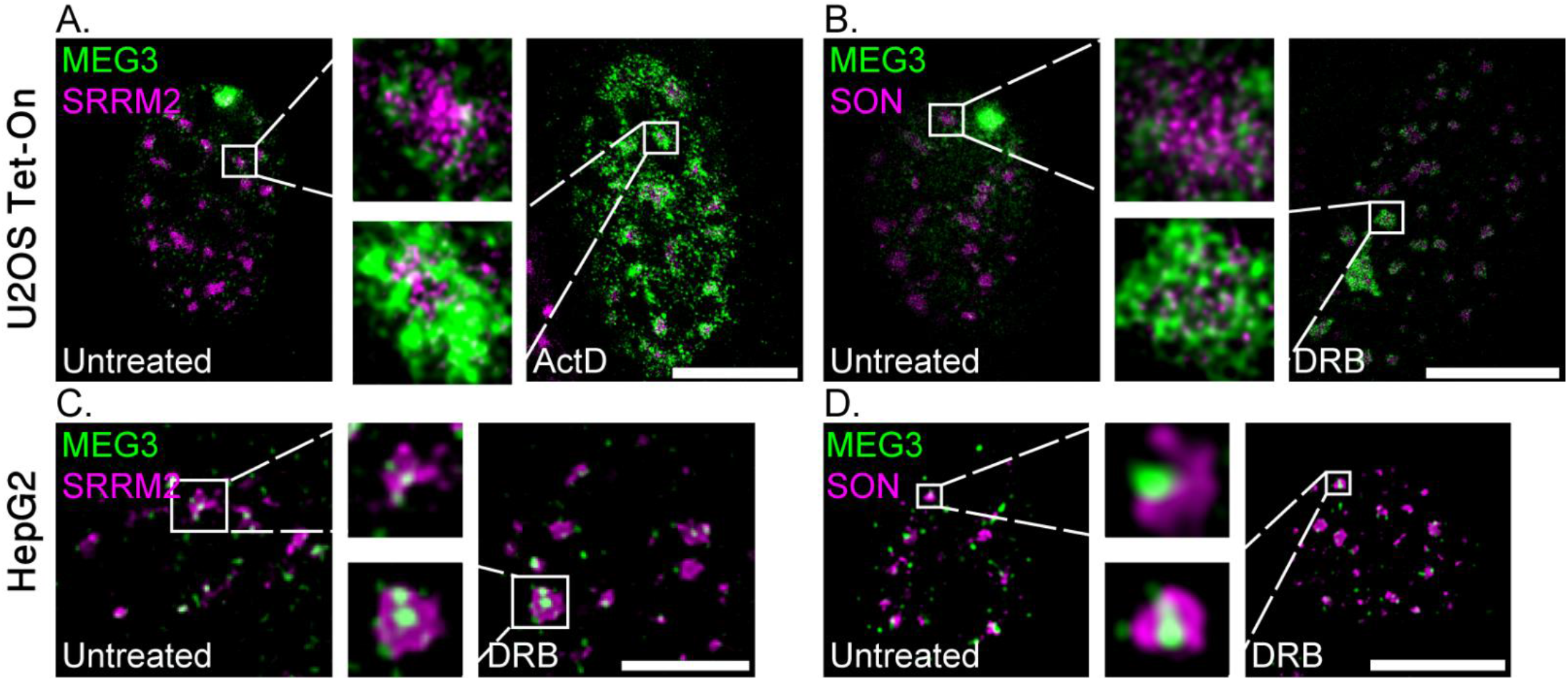
MEG3 can localize to core of the nuclear speckle under transcription inhibition. (**A and B**) STED super-resolution imaging of RNA FISH to MEG3-MS2 RNA (probe to MS2 in green) in U2OS Tet-On cells labeled with (A) SRRM2 or (B) SON staining (magenta), in untreated cells (left) or with ActD/DRB (2 hrs) treatments, respectively (right). Enlargements of boxed areas are in the center. (**C and D**) Confocal imaging of RNA FISH to MEG3 lcRNA in HepG2 cells labeled with (C) SRRM2 or (D) SON (magenta), in untreated cells (left) or with DRB treatment (2 hrs, right). Enlargements of boxed areas are in the center. Bar= 10 μm.

## 4. Discussion

In this study we examined the dynamics of the lncRNA MEG3 within the nucleus of living cells. We found that MEG3 is associated with nuclear speckles under regular conditions and that this association increases when transcription and splicing are inhibited. MEG3 is a lncRNA which plays a crucial role in several key biological processes such as p53 stimulation [66]. It was previously shown that MEG3 transcripts could be found in nuclear speckles where it, among others, colocalizes with the MALAT1 lncRNA [36]. MEG3 is not expressed in all cell types and is expressed at extremely low levels in many cancerous cells. Since it is not expressed in U2OS cells, we could use this cell line for expressing MS2-tagged MEG3 transcripts that can be followed in living cells. The MEG3-MS2 transcript was retained in the nucleus in accordance with endogenous MEG3 behaviour [36]. It partially localized to nuclear speckles similar to the localization of the endogenous MEG3 observed in HepG2 cells. A closer look at the association of MEG3 transcripts with the nuclear speckle marker SRRM2 showed two MEG3 populations, where one appeared to associate with the nuclear speckles while the other was nucleoplasmic. This contrasts with the MALAT1 transcripts which is closely associated with nuclear speckles. The nuclear speckles have subdomains [7] and it appears that the majority of MEG3 transcripts that do associate with the nuclear speckle localize predominantly to the periphery of the nuclear body.

Transcription inhibition causes the nuclear speckles to change structure and become bigger and rounder, since splicing factors are not released from the nuclear speckles into the nucleoplasm under these conditions where there is no splicing [6]. In contrast to the splicing factors that accumulate in nuclear speckles under transcription inhibition conditions, MALAT1 is released from the nuclear speckles and disperses in the nucleoplasm [32]. This is also true for the paraspeckle associated NEAT1 lncRNA [29]. Since MALAT1 and MEG3 commonly both localize to the nuclear speckles and at least partly overlap, we examined whether MEG3 would react to transcription inhibition in a similar manner. However, we found that under transcription inhibition, MEG3 became highly localized at the nuclear speckle. MEG3 therefore differs from MALAT1 and NEAT1 in this respect. Live-cell movies revealed the relatively fast occurrence of this phenomenon. Within 40 minutes of transcription inhibition, the ring formations of MEG3 around the nuclear speckles became prominent. Intensity analysis of MEG3 association with the nuclear speckle periphery indicated increased MEG3 signal at the periphery as well as an increased overlapping with SRRM2 inside the nuclear speckle. Endogenous MEG3 in other cell lines such as hFF and HepG2 behaved similarly to the over-expressed MEG3 in U2OS cells under transcription inhibition. Furthermore, when the cells were treated with the splicing inhibitor PLB, there was increased MEG3 localization to the nuclear speckles. We conclude therefore that the relocalization of MEG3 to nuclear speckles occurs due to the reduced activity of splicing factors in the nucleoplasm and not due to the transcription inhibition.

Nuclear speckles are dynamic structures where the nuclear speckle components shuttle between the nuclear speckle and the nucleoplasm. We wished to understand the nature of the MEG3 nuclear speckle localization and see if the transcripts were anchored to the nuclear speckles or if the association was reversible. Live-cell imaging revealed that by 30 minutes after DRB was removed from the cells, most of the ring formation around the nuclear speckles dispersed in the nucleoplasm in a pattern similar to untreated cells. This time frame of 30 mins correlates with the time required for MALAT1 to re-accumulate in nuclear speckles after release from transcription inhibition [32], suggesting that these two lncRNAs are mutually exclusive with respect to nuclear speckle association. It is possible that MEG3 like other speckle components is stored in or around the nuclear speckle during transcription inhibition perhaps by binding to a certain nuclear speckle component. Another possibility is that MEG3 functions as a carrier for other nuclear speckle components, bringing it to the nuclear speckle during transcription inhibition. The tertiary structure of MEG3 is important for proper function [66]. To examine which regions of the transcript take part in the association with the nuclear speckles we generated a truncated version and found that exons 4 to 8 were not crucial for MEG3 localization to the nuclear speckles. Rather, exons 1 to 3 continued to surround nuclear speckles upon transcription inhibition. It appears that neither core protein SON nor splicing factor SRSF7 or lncRNA binding factor hnRNPK are essential for this localization. Poly(A) RNA signal has been detected in the core the nuclear speckles, however, the only other known poly(A)+ RNA found in the nuclear speckle is MALAT1 lncRNA that localize to the periphery of the nuclear speckle [7]. Super-resolution (STED) microscopy demonstrated that not only is there an increased amount of MEG3 at the nuclear speckle periphery after transcription inhibition, but that the MEG3 transcripts actually integrates into the nuclear speckle core.

Considering that MALAT1, the other known prominent lncRNA at the nuclear speckle, disperses after transcription inhibition, it has not been possible to measure the dynamics of lncRNAs at the nuclear speckle after transcription inhibition. This could be done with MEG3. FRAP measurements showed partial recovery both in treated and untreated cells. However, the recovery was slower for treated cells indicating a slower exchange of the MEG3 transcripts that associate with the nuclear speckle. To examine this on single MEG3 RNPs, we tracked RNPs at the nuclear speckle in living cells and found that during transcription inhibition conditions, the RNPs remained associated with the nuclear speckle periphery. This suggests that the recovery which was seen after the FRAP is due to new MEG3 transcripts joining the nuclear speckles. Altogether, it appears that MEG3 continuously localizes to nuclear speckles during transcription inhibition. Once the MEG3 lncRNA is at the nuclear speckle it appears to remain there.

## 5. Conclusions

Transcription and splicing inhibition cause not only an increased association of splicing factors with nuclear speckles but also the recruitment of the MEG3 lncRNA, in contrast to the MALAT1 lncRNA that is removed from nuclear speckles under these conditions. This shows that relocalization of nuclear factors to nuclear speckles is regulated both on the protein and RNA levels and suggests that MEG3 and MALAT1 lncRNAs are mutually exclusive with respect to nuclear speckle association.

## Author Contributions

Conceptualization, S.E.H. and Y.S-T.; Methodology, S.E.H.; Validation, S.E.H.; Formal Analysis, S.E.H.; Investigation, S.E.H., E.A., M.K.A., A.B., J.G., H.H-L.; Resources, Y.S-T.; Writing - Original Draft Preparation, S.E.H.; Writing - Review & Editing, Y.S-T.; Visualization, S.E.H.; Supervision, Y.S-T.; Project Administration, Y.S-T.; Funding Acquisition, Y.S-T.

## Data Availability Statement

The datasets generated during and/or analysed during the current study are available from the corresponding author on reasonable request.

## Acknowledgments

We thank Irit Shoval, Avi Jacob (BIU Imaging Facility), Hagit Hauschner (BIU FACS Facility) and Jennifer I.C Benichou (BIU) for the assistance with the statistical analysis.

