## Supplemental for "The association of MEG3 lncRNA with nuclear speckles in living cells"

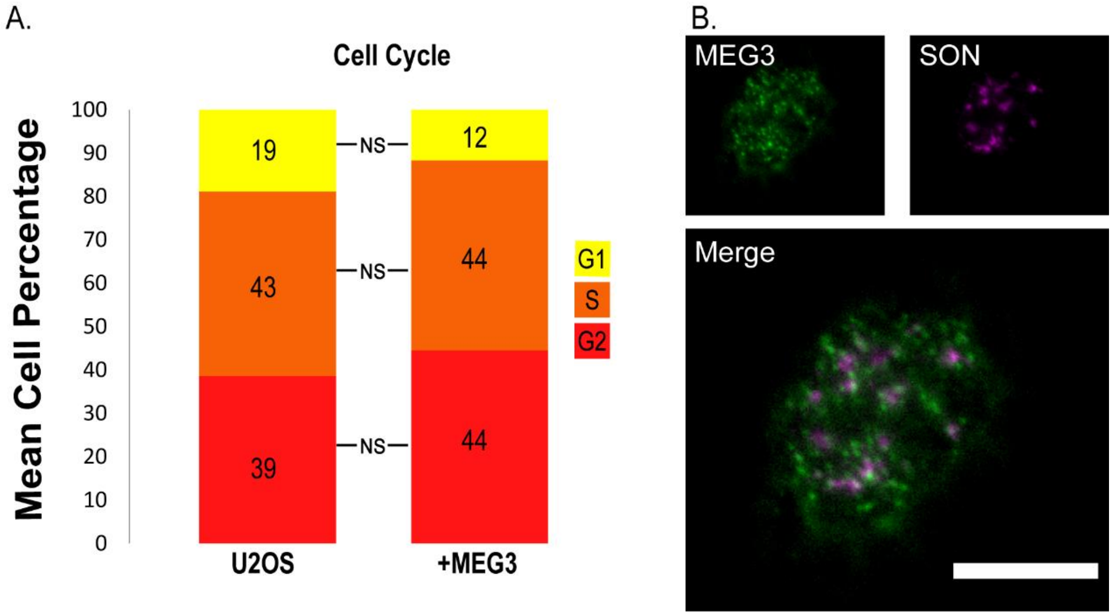


**Supplementary figure S1.** Stable expression of MEG3 in U2OS Tet-On cells did not affect the cell cycle progression of the cells and had a similar expression pattern to endogenous MEG3. (**A**) FACS cell cycle analysis on wildtype U2OS cells and U2OS Tet-On cells expressing MEG3-MS2 transcripts. Percentage of cells found in different stages of cell cycle: G1 (yellow), S (orange), G2 (red) is depicted. The differences between the phases were statistically insignificant (NS) ($p$>0.1). (**B**) HepG2 cells with RNA FISH of endogenous expression of MEG3 transcripts (green) and antibody staining to SON (magenta). Bar= 10 µm.

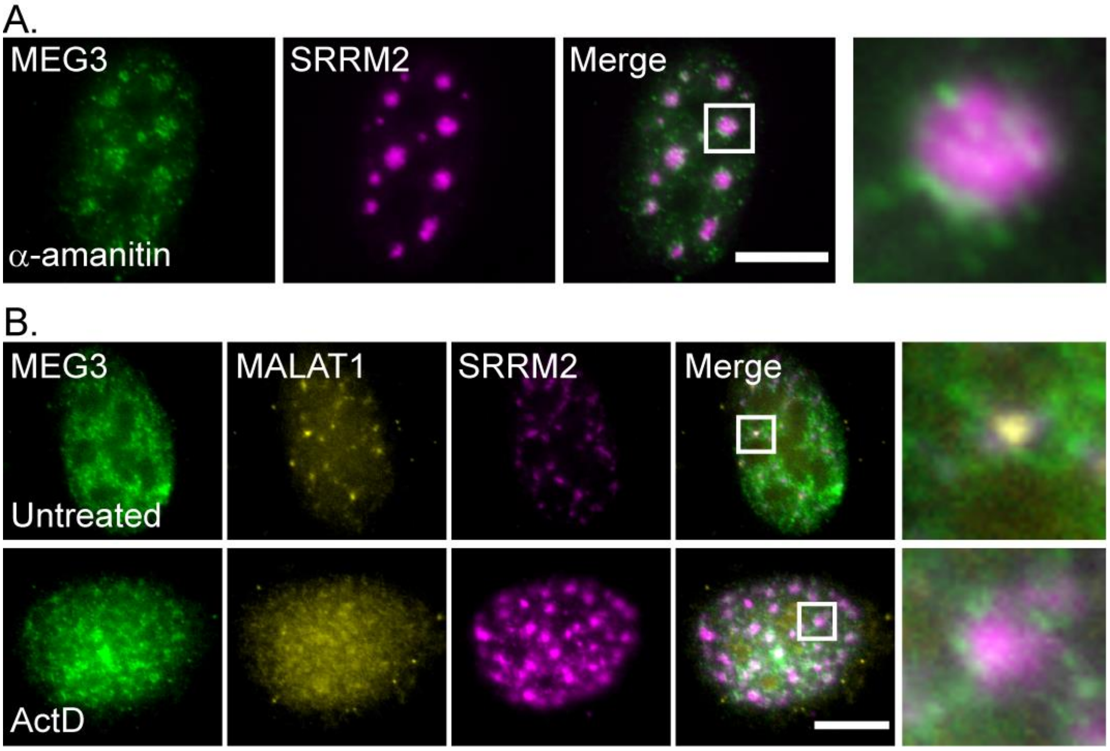


**Supplementary figure S2.** MEG3 relocates to the nuclear speckles after transcription inhibition. (**A**) MEG3 YFP-MS2-CP tagged transcripts (green) and antibody staining of SRRM2 (magenta) transcription inhibition treatment with α-amanitin for 6 hrs before fixation. Enlargements of the boxed regions appear on the right. (**B**) RNA FISH of MEG3-MS2 transcripts with fluorescent probes to the MS2 region (green) in U2OS cells together with endogenous RNA FISH of MALAT1 (yellow) and staining with an antibody to the nuclear speckle marker SON (magenta). (Top) untreated control cells and (bottom) cells treated with ActD for 2 hrs. Bar= 10 µm.

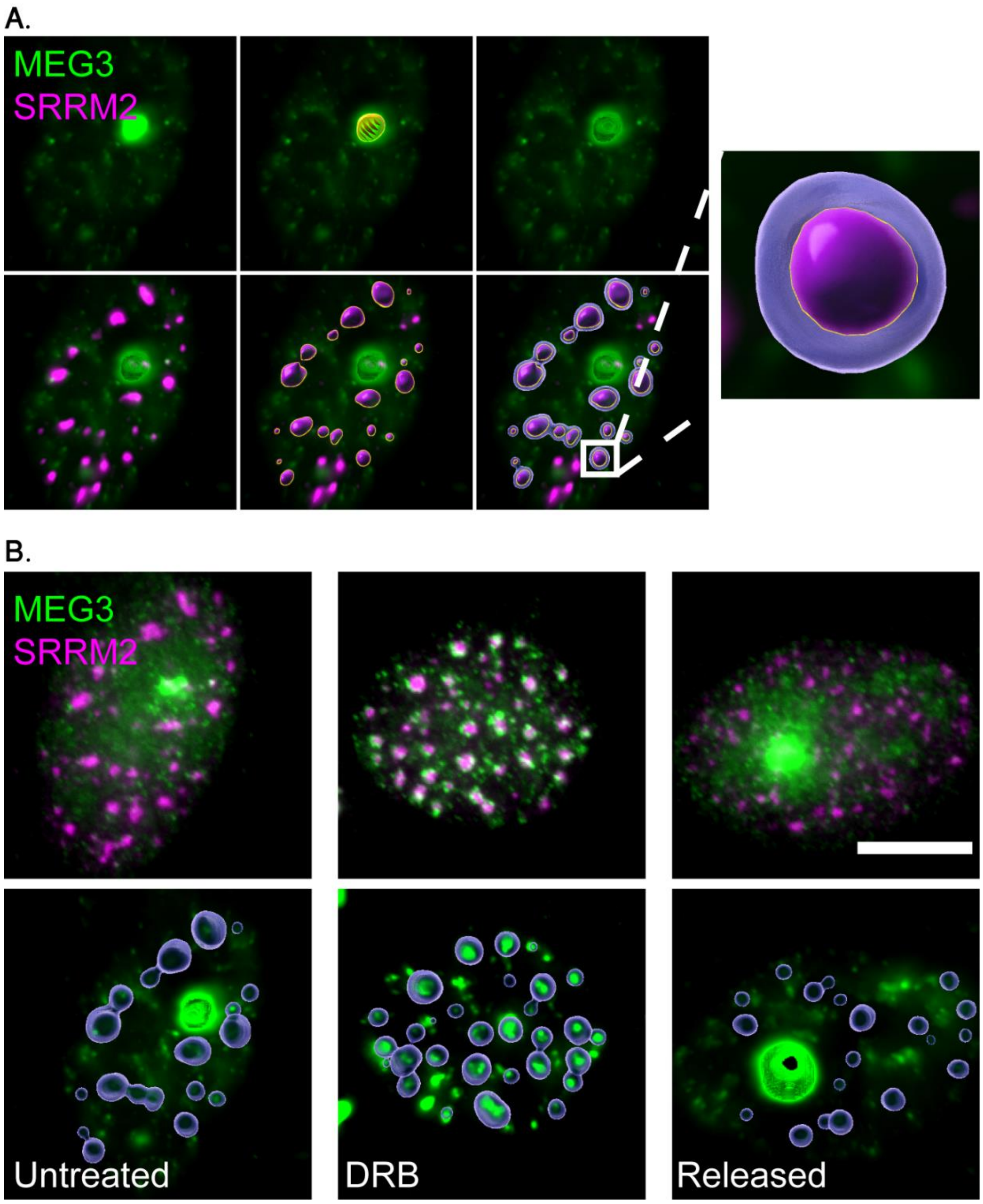


**Supplementary figure S3.** Quantification of MEG3 association with nuclear speckles. Imaris software was used to analyze the MEG3 intensity at the nuclear speckle. (**A**) An untreated cell expressing MEG3 (green) where (top) the transcription site is detected, marked and removed from the analysis. (Bottom) The nuclear speckles (magenta) were then marked and a "bubble" (blue), extending with a threshold value of 0.25 in every direction from the surface of the nuclear speckle, was added. The intensity of MEG3 in the "bubble" were then measured. Enlargement of boxed area is on the right. (**B**) (Top) Original images and (bottom) final Imaris processed images of untreated (left), DRB treated (middle) and DRB-released (right) cells. Bar= 10 μm.

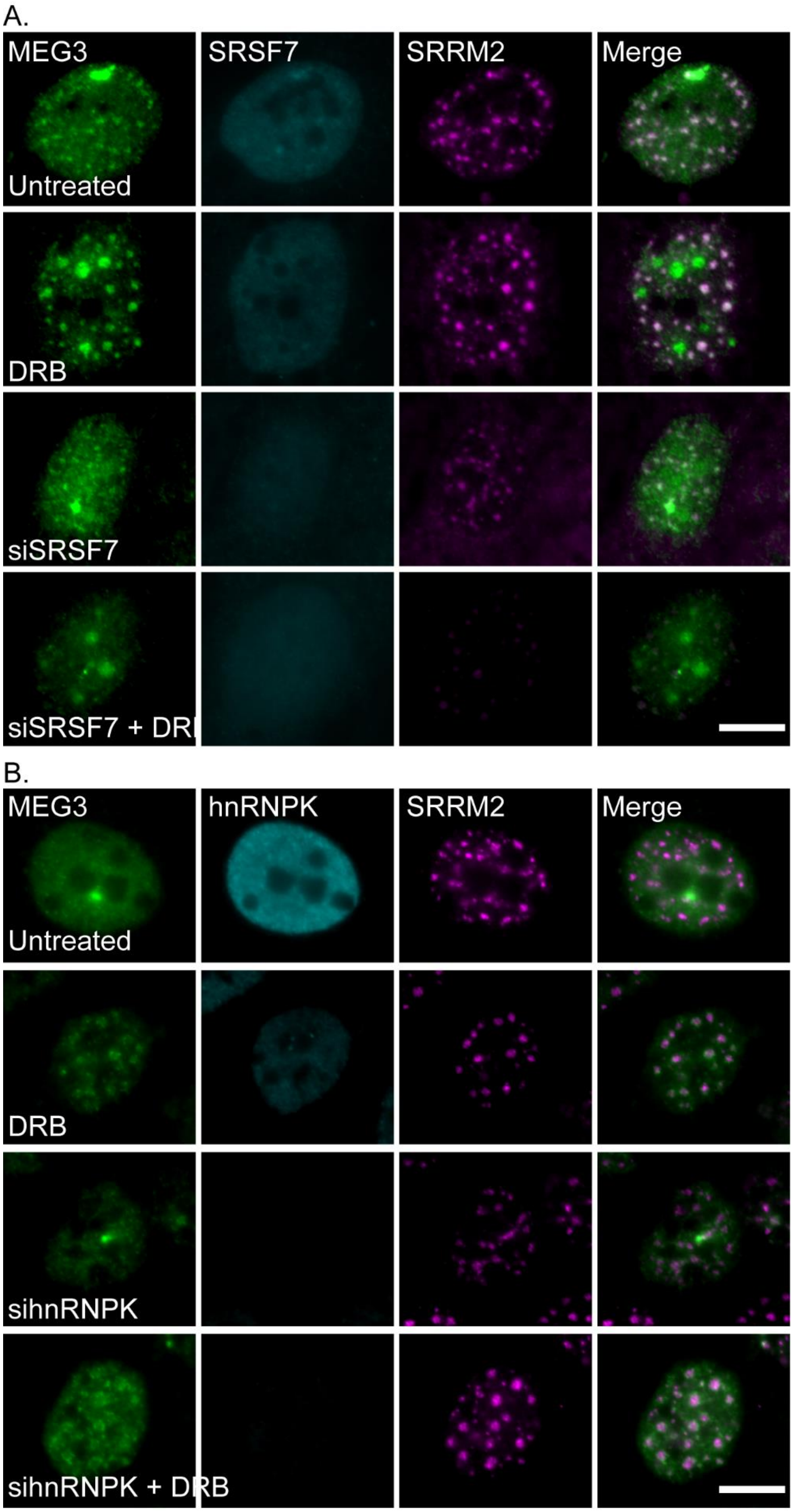


**Supplementary figure S4.** Depletion of splicing factor (**A**) SRSF7 or (**B**) hnRNKP proteins does not change MEG3 association with nuclear speckles. Knockdown of SRSF7/hnRNPK (cyan) in cells labeled by RNA FISH for MEG3-MS2 transcripts (green) and staining SRRM2 (magenta). (Top) Untreated control cells, (top middle) cells treated with DRB for 2 hrs, (bottom middle) siRNA knockdown, (bottom) siRNA knockdown with DRB treatment for 2 hrs. Bar= 10 µm.

**Supplemental Movie Legends**

**Supplementary Movie S1.** Live-cell imaging of U2OS cells stably expressing MEG3-MS2 tagged with YFP-MS2-CP after Dox activation (MEG3 grey signal is the inverted version of the original images). Small dots are MEG lncRNPs and the large dot is the actively transcribing gene. There is a gradual increase of MEG3 signal over a 3 hr period. Imaged every 10 min for 3 hrs.

**Supplementary Movie S2.** Live-cell imaging of U2OS cells stably expressing MEG3-MS2 tagged with YFP-MS2-CP (MEG3 grey signal is the inverted version of the original images) where cells were treated with DRB 15 min before the start of the movie. Nuclear speckle association can be observed after about 40 min and remains throughout the movie. Imaged every 5 min for 1 hr and 40 min.

**Supplementary Movie S3.** Live-cell imaging of U2OS cells stably expressing MEG3-MS2 tagged with YFP-MS2-CP (MEG3 grey signal is the inverted version of the original images). The cells were treated with DRB for 2 hrs followed by release from the DRB immediately prior to the imaging. MEG3 is at the onset localized to the nuclear speckles and gradually disperses in the nucleoplasm. There is a complete loss of ring formation after 30 min of release from DRB. Imaged every 5 min for 1 hr and 20 min.

**Supplementary Movie S4.** FRAP live-cell imaging of an untreated U2OS cell stably expressing (left) MEG3-MS2 tagged with YFP-MS2-CP (yellow) and over expressing (right) Cerulean-SRSF3 (cyan). A red ring indicates the area bleached. MEG3 shows a clear nuclear speckle localization at the onset of the imaging and then there is recovery of the YFP signal at the specific nuclear speckle where the MEG3 signal was bleached.
